# Characterizing Low-Frequency Artifacts During Transcranial Temporal Interference Stimulation (tTIS)

**DOI:** 10.1101/2022.03.07.483213

**Authors:** Jill von Conta, Florian H. Kasten, Branislava Ćurčić-Blake, Klaus Schellhorn, Christoph S. Herrmann

## Abstract

Transcranial alternating current stimulation (tACS) is a well-established brain stimulation technique to modulate human brain oscillations. However, due to the strong electro-magnetic artifacts induced by the stimulation current, the simultaneous measurement of tACS effects during neurophysiological recordings in humans is challenging. Recently, transcranial temporal interference stimulation (tTIS) has been introduced to stimulate neurons at depth non-invasively. During tTIS, two high-frequency sine waves are applied, that interfere inside the brain, resulting in amplitude modulated waveforms at the target frequency. Given appropriate hardware, we show that neurophysiological data during tTIS may be acquired without stimulation artifacts at low-frequencies. However, data must be inspected carefully for possible low-frequency artifacts. Our results may help to design experimental setups to record brain activity during tTIS, which may foster our understanding of its underlying mechanisms.

## 1 Introduction

Transcranial alternating current stimulation (tACS) is a well-established brain stimulation technique to modulate brain oscillations and human behavior (Herrmann, Rach, Neuling, & Strüber, 2013; Kanai, Chaieb, Antal, Walsh, & Paulus, 2008; Kasten & Herrmann, 2017; Vossen, Gross, & Thut, 2015; Zaehle, Rach, & Herrmann, 2010). However, the simultaneous measurement of tACS effects during neurophysiological recordings in human electroencephalography (EEG) and magnetoencephalography (MEG) is challenging due to the strong electro-magnetic artifacts induced by the stimulation current. Those artifacts spectrally overlap with the brain oscillation under investigation and thereby hinder direct insights on the effects of tACS during stimulation (Kasten & Herrmann, 2019). Several analysis approaches, such as template subtraction (Helfrich et al., 2014), or spatial filtering (Kasten, Maess, & Herrmann, 2018; Neuling et al., 2015) aimed to suppress the stimulation artifact in M/EEG data. The complete restauration of the intrinsic brain activity remains, however, challenging (Noury, Hipp, & Siegel, 2016; Noury & Siegel, 2018).

Recently, amplitude modulated tACS (AM-tACS) has been suggested as an approach to overcome the stimulation artifact at the target frequency (Witkowski et al., 2016). During AM-tACS a high-frequency carrier sine wave is modulated at the target frequency. In theory, AM-tACS exhibits power at the carrier frequency and two side bands (carrier frequency ± modulation frequency), circumventing the artifact at the modulation frequency. Based on simulations, Kasten, Negahbani, Fröhlich, & Herrmann (2018) investigated low-frequency artifacts caused by the equipment in electrophysiological measurements during AM-tACS. They showed that non-linear properties of stimulation and recording hardware cause small changes to the AM-waveform which reintroduce low-frequency artifacts at the frequency of the amplitude modulation. If stimulation or recording hardware operate in a non-linear fashion, the output signal of a device is not a linear product of the (e.g. amplified) input signal. Further, the authors showed that low-frequency artifacts were present in phantom- and human-measurements in different experimental setups during AM-tACS.

Transcranial temporal interference stimulation (tTIS) has recently been proposed as a possible non-invasive technique for selectively stimulating deep brain regions (Grossman et al., 2017; Wessel et al., 2021). During tTIS two high-frequency sinusoidal currents of slightly different frequencies are applied to the brain and interfere in regions of overlap. The resulting sum signal features a signal with a fluctuating envelope depending on the difference of the two high-frequency sinusoids. For AM-tACS, the amplitude modulation is generated by multiplication of the high and low frequency sinusoids before the signal is forwarded to the signal generators (Digital-Analog-Converter (DAC) and the constant current source of the stimulator). For tTIS, the quasi-amplitude modulation (i.e. the expression of the envelope or the beat rhythm at the difference frequency) emerges inside the brain, after the independent generation of two pure high-frequency sine-waves (independently generated).

The aim of the current study is to characterize low-frequency artifacts due to tTIS in neurophysiological recording methods. Due to the generation of the signal (two pure sine-waves are generated independently), tTIS circumvents non-linearities inherent in the signal generation hardware. Therefore, we hypothesized that low-frequency artifacts due to tTIS could be absent or, at least, smaller in size, if we measure with a recording device (such as M/EEG), characterized by high linearity. In order to test this hypothesis, we simulated the resulting signal for tTIS, accounting for non-linearities of the signal generator- and recording-hardware. Moreover, we systematically tested three different recording devices while stimulating a phantom head with tTIS.

## 2 Methods

### 2.1 Simulation of non-linear signal generation and acquisition

To assess the role of non-linear hardware components for low-frequency artifacts during tTIS, we adapted the simulation regarding the influence of non-linear input-output function of a system presented in Kasten et al. (2018). Specifically, we extended the simulation by dissociating the influence of the hardware at the signal generation stage, as well as at the signal recording stage to the stimulation signal. We contrasted the influence of non-linearities (inherent in the signal generation- and recording-hardware) for an amplitude modulated signal (AM-tACS), against the influence of non-linearities for two independently generated sine waves, interfering inside the brain to an amplitude modulated signal (tTIS).

Non-linear input/output properties of the signal generation and recording stages were modelled using 6^th^ degree polynomial functions of the form:

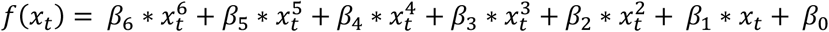

where *x* is the amplitude of the input signal at time *t* and β is a weight for the non-linear term. Weights for the function were approximated by using values previously observed for a complete tACS signal generation and EEG acquisition circuit in our previous work (Kasten et al. 2018). Signals were fed through the function twice to simulate a signal generation and a signal recording stage. For the AM-waveform the signal was created prior to both stages using the following formula:

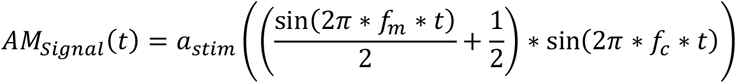

where *a_stim_* is the stimulation amplitude, *f_m_* is the modulation frequency and *f_c_* is the carrier frequency. The resulting signal is an AM-waveform with 50% modulation depth. In contrast, for the tTIS signal, two pure sinewaves with frequencies of f_1_ = 195 Hz and f_2_ = 205 Hz were created. The two sinewaves were separately fed through the polynomial function simulating the signal generation stage and then linearly combined. Subsequently, the resulting tTIS signal, which now exhibits an amplitude modulation at a frequency equal to the difference of f_1_ and f_2_, was fed through the polynomial function a second time, to simulate the signal acquisition stage.

### 2.2 Phantom M/EEG recordings

In addition to our simulations, we systematically tested three different recording devices while stimulating a phantom head (melon) with tTIS: (1) an MEG, (2) a passive- and (3) an active-EEG system. Specifically, we recorded MEG with a 306-channel whole head MEG system by Elekta Neuromag Triux (Elekta Oy, Helsinki, Finland), sampled at 5 kHz. Moreover, we measured with two different EEG systems: a passive- (16-bit amplifier, BrainAmp system, sampled at 5 kHz) and an active- (24-bit amplifier, actiChamp System, sampled at 10 kHz) EEG system, both manufactured by Brain Products GmbH (Gilching, Germany) with (active) Ag/AgCl electrodes attached to the phantom head at positions roughly corresponding to a nose (reference electrode), forehead (ground electrode) and Pz montage (positions were approximated), while keeping the impedances of the EEG electrodes below 10 kΩ. While measuring each recording device (MEG, passive- and active EEG), we systematically stimulated with two different carrier frequencies (500 Hz and 1000 Hz), as well as with several different modulation frequencies (10 Hz, 11 Hz, and 23 Hz). For an overview across the recording setups see *Table 1*. The stimulation signal was digitally generated at a rate of 100 kHz, transferred to a digital analog converter (NI-USB 6251, National Instruments, Austin, TX, USA), which was connected to optically isolated remote inputs of two battery-operated constant current stimulators (Advanced DC STIMULATOR PLUS, neuroConn, Ilmenau, Germany). The impedances of the stimulation electrodes were kept below 10 kΩ. The stimulation intensity was 1 mA for MEG- and active EEG-recordings. Due to limits of the recording hardware (lower amplitude rang for 16-bit digitalization), the stimulation intensity was 0.1 mA for passive EEG recordings.

**Table 1.** Recording Setups.

|  | (1) MEG | (2) Passive EEG | (3) Active EEG |
| --- | --- | --- | --- |
| System | Elekta Neuromag Triux | brainAmp system, 16-bit amplifier | actiChamp System, 24-bit amplifier |
| Sampling Rate | 5 kHz | 5 kHz | 10 kHz |
| Carrier Frequency | 500 Hz<br>1000 Hz | 500 Hz<br>1000 Hz | 500 Hz<br>1000 Hz |
| Modulation Frequency | 10 Hz<br>11 Hz<br>23 Hz | 10 Hz<br>11 Hz<br>23 Hz | 10 Hz<br>11 Hz<br>23 Hz |
| Intensity | 1mA | 0.1mA | 1mA |

### 2.3 Data Analyses

To assess the frequency content of the simulated stimulation waveforms at different stages of the signal generation-recording circuit, power spectra were computed using Fast Fourier Transforms. Simulations were computed for 1 sec segments of data, resulting in a frequency resolution of 1 Hz. Spectra were assessed for the original waveforms (before being fed through any non-linear function), after the signal generation stage, and after the signal acquisition stage.

M/EEG data were analyzed using Matlab 2018b (The Mathworks Inc., Natick, MA, USA) and the Fieldtrip toolbox. Specifically, M/EEG data were high pass filtered at 1 Hz, and the signals were cut into 1-sec epochs. On each of these segments, FFTs (4-sec zero-padding, Hanning window) were computed and averaged. For the MEG data, results are shown for an exemplary magnetometer channel (*MEG0732*).

## 3 Results

The simulation suggests, that if the AM-waveform is created directly in the stimulator (as for AM-tACS), we clearly see low-frequency artifacts at the difference frequency after the signal generation and the signal recording stage (see Figure 1A, *top row*). If, however, we send multiple pure sine waves through the system (as for tTIS), no low-frequency artifact can be found in the spectrum after the signal generation stage (see Figure 1A, *bottom row*). This shows a clear advantage of tTIS over traditional AM-tACS. Nonetheless, if the resulting AM-wave is then sampled by another non-linear recording device (e.g. M/EEG recording device), the low-frequency artifact is reintroduced to the recorded signal (see Figure 1A, *bottom row*). For MEG and the passive EEG system, we can clearly see the low-frequency artifact in the data at the different modulation frequencies (10 Hz, 11 Hz, and 23 Hz) for both carrier frequencies (500 Hz and 1000 Hz) (see Figure 1B left and middle, zoomed spectra 1-40 Hz). In contrast, when recording with the active EEG system, the low-frequency artifacts at the different modulation frequencies are not visible in the data for neither one of the carrier frequencies (see Figure 1B right).

**Figure 1.**
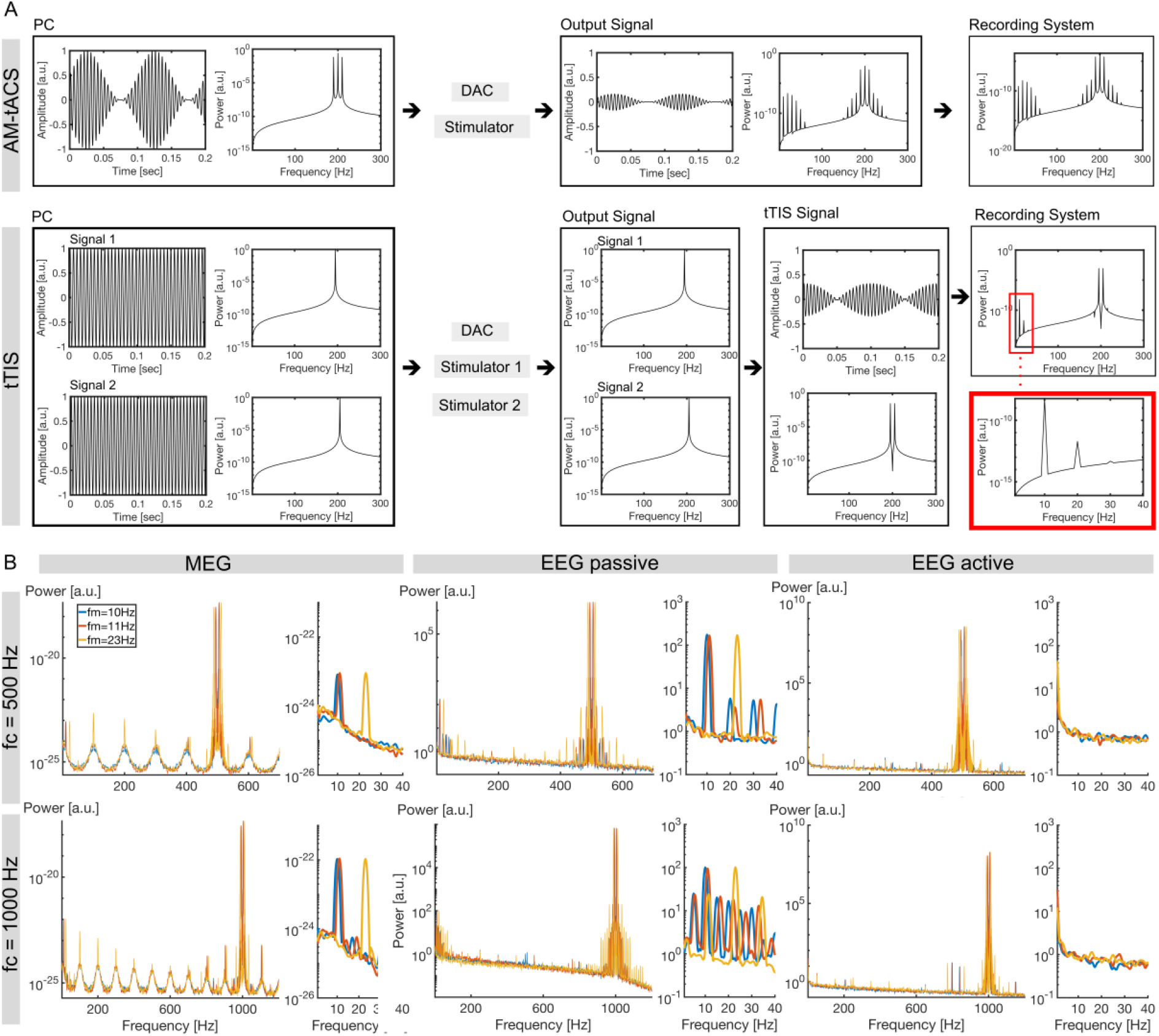
Results. A. Simulation results. For AM-tACS, the amplitude modulated signal is generated (carrier frequency of 200 Hz, modulation frequency of 10 Hz) within the computer, characterized by power at the carrier frequency and two side bands (carrier frequency ± modulation frequency). The signal is forwarded to the simulation of the signal generators (DAC and stimulator). The output signal shows that the power spectra of the amplitude modulated signal exhibits low frequency artifacts at the modulation frequency, and harmonics. After simulating a non-linear measurement device, we can clearly see the low frequency artifacts in the spectra (top row). For tTIS, two sinusoidal signals (Signal 1 oscillates at 195Hz, Signal 2 oscillates at 205Hz) are generated and forwarded to the simulation of the signal generators (DAC and stimulators). The output signals of both single sinusoids show one peak at the stimulation frequency (195 Hz and 205 Hz). Adding the sinusoids to the amplitude modulated signal, we can see no low-frequency artifact in the signal. We solely can see the peaks at the initial frequencies of both sinusoids (195Hz and 205 Hz). If, however, we simulate a non-linear measurement device, we again can see low-frequency artifacts in the data (at the modulation frequency and its harmonics as indicated in the red boxes). B. Recording results for MEG, passive EEG, and active EEG. The low-frequency artifact is clearly visible at different modulation frequencies (f_m_) for MEG (visualized for one gradiometer channel), and passive EEG (left and middle). In contrast, no low-frequency artifact is visible in active EEG recordings (right). Note that all frequency spectra are displayed in log scale.

## 4 Discussion

Our simulation confirms that during tTIS we avoid artifacts caused by the non-linearity of the stimulators (due to the nature of its signal generation), but the non-linearity of the recording system is crucial for the occurrence of low-frequency distortions during tTIS. This indicates that low-frequency artifacts in the recorded signal is mainly caused by non-linearities inherent in the recording hardware (not in the signal generation hardware). Indeed, experimental data show that the artifact arises in the recorded signal due to non-linearities inherent in the recording hardware for MEG, and the passive EEG system. In contrast, data recorded with the active EEG system show no low-frequency artifact. Note that the active EEG system has a larger dynamic range and, therefore, has a bigger range where the device can sample linearly. The results of the current investigation suggest that it is possible to record human brain activity during tTIS without a low-frequency artifact present in the data masking the brain activity at frequencies that are in the range of regular neural frequencies. However, the quality of the neurophysiological data during tTIS highly depends on the recording hardware that is used. As reported for the MEG- and the passive EEG-system, the artifact at the modulation frequency can be easily introduced to the data on the recording level. However, if the measurement is done using an almost linear recording device (such as the active EEG system in the current investigation), the artifact at the modulation frequency can be minimized making the signal useful for investigation of brain functions. Note, the current investigation is limited to phantom data with specific stimulation parameter (stimulation intensity, frequencies). Therefore, future neurophysiological studies with simultaneous tTIS must be inspected carefully in regard to possible low-frequency artifacts. Overall, it has to be noted that the non-linearities of the used recording devices are extremely small and negligible in regular recordings of human brain activity. However, they do become relevant when additional signals at high intensities such as during transcranial stimulation are introduced to the recordings, which is arguable not the main purpose these devices were designed for.

The results of the current investigation suggest that it is feasible to simultaneously stimulate using tTIS and record good quality data without a low-frequency artifact, given a linear recording device. While tTIS is a novel method and little is known about its efficacy in humans, our work may help to design recording setups that allow recording of brain activity during stimulation and foster our understanding of its underlying mechanisms.

## Acknowledgements

This research was supported by the Neuroimaging Unit of the Carl von Ossietzky University Oldenburg funded by grants of from the German Research Foundation (3T MRI INST 184/152-1 FUGG and MEG INST 184/148-1 FUGG). This work was funded by the Deutsche Forschungsgemeinschaft (DFG, German Research Foundation) under Germany’s Excellence Strategy – EXC 2177/1 – Project ID 390895286.

## Author contributions

JC, FHK, BCB, KS and CSH conceived the study. JC analyzed the data. JC, FHK, BCB, KS and CSH wrote the manuscript.

